# Paralog Explorer: a resource for mining information about paralogs in common research organisms

**DOI:** 10.1101/2022.07.22.501126

**Authors:** Yanhui Hu, Ben Ewen-Campen, Aram Comjean, Jonathan Rodiger, Stephanie E. Mohr, Norbert Perrimon

## Abstract

Paralogs are genes which arose via gene duplication, and when such paralogs retain overlapping or redundant function, this poses a challenge to functional genetics research. Recent technological advancements have made it possible to systematically probe gene function for redundant genes using dual or multiplex gene perturbation, and there is a need for a simple bioinformatic tool to identify putative paralogs of a gene(s) of interest. We have developed Paralog Explorer (https://www.flyrnai.org/tools/paralogs/), an online resource that allows researchers to quickly and accurately identify candidate paralogous genes in the genomes of the model organisms *D. melanogaster, C. elegans, D. rerio, M. musculus*, and *H. sapiens*. Paralog Explorer deploys an effective between-species ortholog prediction software, DIOPT, to analyze within-species paralogs. Paralog Explorer allows users to identify candidate paralogs, and to navigate relevant databases regarding gene co-expression, protein-protein and genetic interaction, as well as gene ontology and phenotype annotations. Altogether, this tool extends the value of current ortholog prediction resources by providing sophisticated features useful for identification and study of paralogous genes.

## Introduction

Genes that arise as a result of gene duplication are known as paralogs, and in cases where they retain overlapping function, it can represent a particular challenge to functional analysis [1]. Specifically, while loss-of-function experiments have been enormously successful to characterize individual gene function, this approach can fail when a target gene has a redundant or partially redundant paralog that can compensate in its absence. For example, large-scale studies in yeast provide evidence that, in aggregate, knocking out singleton genes (those without any paralog) tends to produce a stronger phenotypic effect than knocking out one member of a paralog pair, likely due at least partially to paralog-based redundancy [2]. Similarly, gene essentiality studies across nearly 600 human cancer cell lines revealed that paralogs were far less likely than singleton genes to be essential [3].

Importantly, paralogs are not a minor curiosity in eukaryotic genomes. In fact, gene duplication and functional divergence has long been recognized as one of the most fundamental and widespread sources of evolutionary novelty [4], and paralogs are extraordinarily widespread in genomes. In the human genome, for instance, 70.5% of all genes are estimated to have at least one paralog [5]. Similarly, 67% of *Drosophila* genes have been estimated to have at least one paralog [1]. While many paralogs have diverged functionally and encode proteins of unrelated or non-overlapping roles, there is undoubtedly a large amount of functional diversity yet to be characterized which has thus far remained invisible to standard gene loss-of-function studies.

In recent years, it has become increasingly possible to perform double-or multiplex loss-of-function experiments using scalable techniques as double RNAi or CRISPR-based techniques. To date, massively parallel double-knock CRISPR approaches have been primarily applied to cell culture experiments, where it is possible to introduce dual-sgRNA libraries targeting tens of thousands of gene pairs [6-9]. However, in vivo CRISPR-based techniques for dual- and multiplex loss-of-function experiments are rapidly being developed for model organisms [10-14].

To facilitate the functional studies of paralogs in model organisms, both in cell culture and in vivo, we have developed a simple but effective bioinformatic tool, Paralog Explorer (https://www.flyrnai.org/tools/paralogs/web/) to identify and explore paralogous genes. Paralog Explorer allows users retrieve paralogs based on a single or multi-gene query, across a wide range of sequence similarity, and to provide relevant comparative information about the retrieved paralog pairs. Paralog Explorer is based on *Drosophila* RNAi Screening Center Integrative Ortholog Prediction Tool (DIOPT), which was developed to identify orthologous genes between species using an integrative approach. By focusing the DIOPT algorithm within, as opposed to between, species, Paralog Explorer identifies paralogs within a given genome. Further, the resource retrieves associated public data and annotations such as chromosomal location, sequence similarity, expression data from different tissues, and protein-protein or genetic interactors. This information can help researchers predict which paralogous genes might act redundantly or otherwise in concert with one another, and thus to assist in designing targeted small-or large-scale experimental studies.

## Materials and Methods

### Paralog information

Paralog information was obtained from DIOPT database release 8 [15]. DIOPT integrates 17 existing algorithms and resources and use a simple voting system for rapid identification of orthologs and paralogs among major model organisms. The organisms included in the Paralog Explorer resource are the nematode worm *C. elegans*, the fruit fly *D. melanogaster*, the mouse *M. musculus*, the zebrafish *D. rerio*, and human *H. sapiens*. Protein alignment information, including alignment length, percent similarity, and percent identity, were also imported from DIOPT. In addition, for genes in each paralog pair, the orthologs in more ancient species such as yeast orthologs for human, mouse, zebrafish, worm and Drosophila paralogous genes, and Drosophila orthologs for paralog genes in mammals are also analyzed. The common orthologs shared by both genes in a pair are identified and stored in database for display. Data files were exported from DIOPT in text format and were further processed using a local program. The output files are uploaded into a mySQL database.

### Integration of omics datasets

For each gene in a paralog pair, we retrieved and integrated protein-protein interaction and genetic interaction data from MIST [16]. In addition, we also identified interactors in common for each paralog pair. Tissue-or developmental stage-specific expression datasets were also integrated. For each Drosophila paralog pair, modENCODE tissue-, developmental stage-, cell line-, and treatment-specific expression profiles provided by FlyBase were integrated [17-20]. In addition, Pearson correlation coefficient scores were calculated for each dataset and synexpression analysis was done. For human paralog pairs, tissue-specific expression data from GTEx Portal (https://gtexportal.org/home/) were integrated and synexpression analysis was done [21].

### Integration of other annotation

The ‘slim’ versions of gene ontology (GO) annotations were retrieved from NCBI. For each gene pair, common GO slim terms were identified and stored. Genome coordinates were retrieved from NCBI EntrezGene. Phenotype annotations from FlyBase (r6.45) and gene group annotations from GLAD [22] were retrieved. This type of information is subject to update periodically in Paralog Explorer.

### Web-based tool development

The Paralog Explorer web tool (https://www.flyrnai.org/tools/paralogs/) can be accessed directly or found at the ‘Tools Overview’ page at the DRSC/TRiP Functional Genomics Resources website (https://fgr.hms.harvard.edu/tools). The backend was written in PHP using the Symfony framework and the front-end HTML pages take advantage of the Twig template engine. The JQuery JavaScript library with the DataTables plugin is used for handling Ajax calls and displaying table views. The Bootstrap framework and some custom CSS are used on the user interface. A mySQL database is used to store the integrated information and analysis results (e.g., Pearson correlation co-efficient scores for synexpression). Both the website and databases are hosted on the O2 high-performance computing cluster, which is made available by the Research Computing group at Harvard Medical School.

## Results

### Database content and user interface features

To build Paralog Explorer, we retrieved all paralog predictions from DIOPT for human, mouse, zebrafish, fly and worm of DIOPT score, equal or larger than 2 (Figure 1). The ‘DIOPT score’ is the number of algorithms (eg. 7 out of 16 for human and *Drosophila* ortholog mapping) that support a given prediction, which we previously showed provides a measure of confidence in each prediction [15]. Protein alignment information, including the alignment length, percent similarity, and percent identity, was also imported from DIOPT. We find that 34%-69% of paralog pairs in Paralog Explorer are supported by 4 or more algorithms and 15-39% have score equal or more than 6 (Table 1). We also imported Gene Ontology (GO) terms [23, 24], protein-protein and genetic interaction data from MIST [16], expression data from publicly-available databases such as modENCODE [18] and GTEx [21], and phenotype data for *Drosophila* from FlyBase [25] (Figure 1).

**Table 1.** Summary of predicted paralogs in Paralog Explorer.

| **NCBI TaxID**<br>**(organism)** | **Maximum**<br>**DIOPT score** | **Gene count**<br>**(DIOPT score>=2)** | **Gene count**<br>**(DIOPT score>=4)** | **Gene count**<br>**(DIOPT score>=6)** |
| --- | --- | --- | --- | --- |
| 6239 (worm) | 10 | 15333 | 5206 (34%) | 2245 (15%) |
| 7227 (fly) | 10 | 10092 | 4776 (47%) | 2486 (25%) |
| 7955 (zebrafish) | 9 | 19788 | 13604 (69%) | 7685 (39%) |
| 9606 (human) | 10 | 18114 | 11153 (62%) | 6559 (36%) |

**Figure 1.**
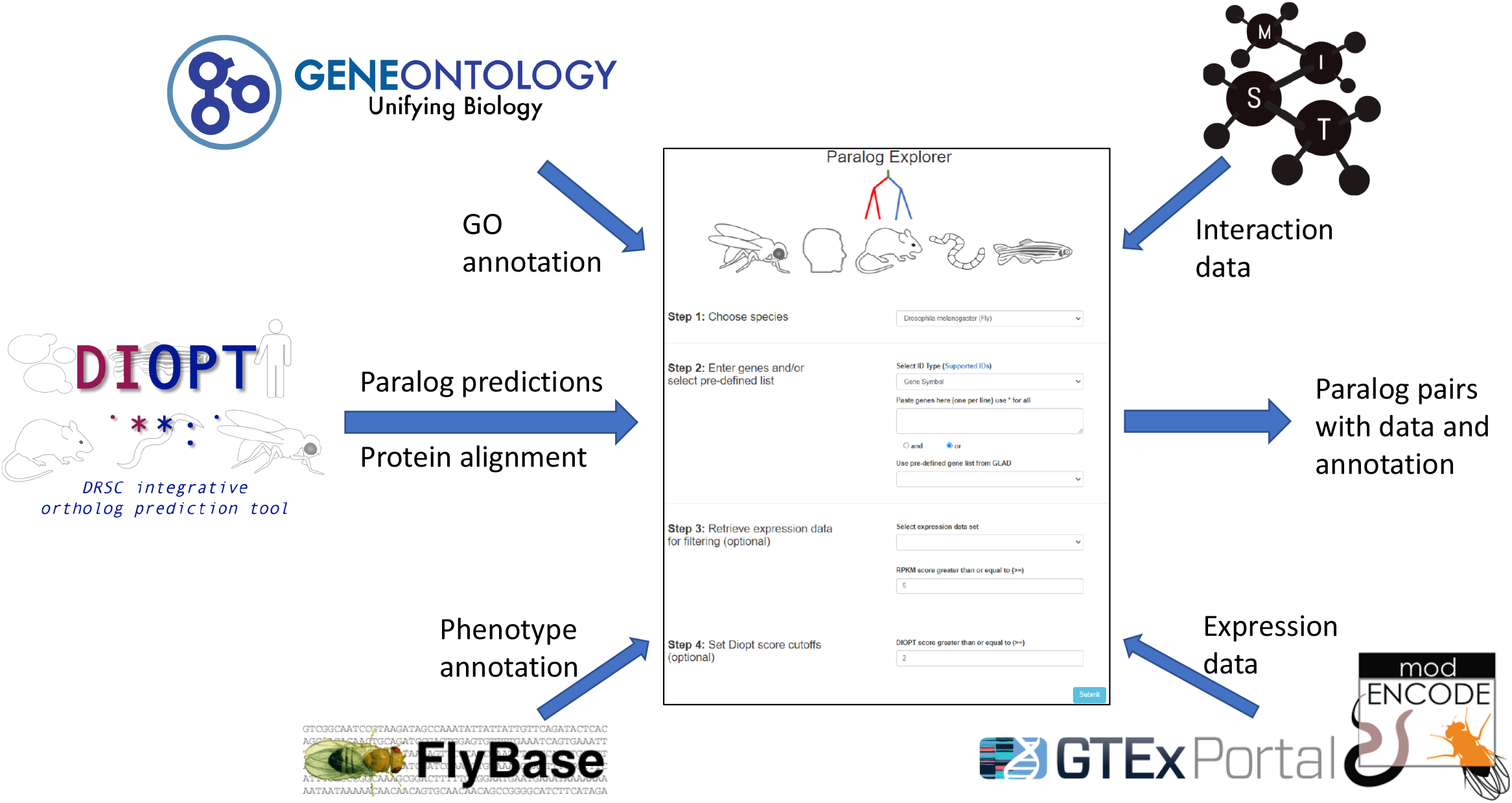
Sources for information included in Paralog Explorer. Paralog Explorer was built based on the integration of paralog predictions from DIOPT, PPI and genetic-interaction data from MIST, expression data from GTEx and modENCODE as well as gene ontology and phenotype annotation from GO consortium and FlyBase, respectively.

With the Paralog Explorer web-tool, users can query a specific gene of interest, a list of genes, or any one of several pre-computed gene lists from GLAD [22]. In addition, users can establish a filter based on DIOPT score, and for *Drosophila* and human genes, can establish a cut-off of transcriptional expression level in transcriptomic datasets from various tissues for both genes in a pair. Altogether, the user interface is designed to allow users to answer a variety of questions. These include very straightforward questions such as, does my gene of interest have one or more paralogs? Or, which of the genes in a list (e.g. hits from a genetic screen) have paralogs? Paralog Explorer also supports more complex queries such as, what are all the paralogous genes expressed in a given tissue? What are all the paralogous genes encoding transporters that are expressed at high levels in the adult digestive system?

For each query, the Paralog Explorer web-tool reports the total number of paralogs identified within a genome, each of which is shown on a separate line. For each paralog pair, Paralog Explorer displays information including the DIOPT score of the paralog, the genomic location of each gene, as well as various measures of protein alignment and Gene Ontology (GO) annotation for each member of the paralog pair, as well those GO terms common to both paralogs.

For each gene in a paralog pair, Paralog Explorer also reports the top-scoring ortholog(s) from the distantly-related outgroup yeast (*S. cerevisiae*), if such orthologs exist. This allows users to assess whether both paralogs in an animal model correspond to a single ortholog in yeast, which may assist in generating functional hypothesis. Similarly, for all non-*Drosophila* organisms, Paralog Explorer returns the closest fly ortholog for each member of a paralog pair. For example, PTPN11 and PTPN6, a paralogous gene-pair in humans, are both orthologous to *csw* in the *Drosophila* genome. This information can help to clarify whether a given paralog pair is the result of a lineage-specific gene duplication, or whether the duplication predated the divergence of these lineages [26].

The tool also integrates several -omics datasets of protein-protein and genetic interaction, to identify genetic and physical interactors of each gene in the paralog pair. After gene duplication, protein interactions can be conserved among paralogs [27]. Thus, paralogous genes that share common partners may be more likely to be functionally related and the interaction data can help user to prioritize the gene pairs when design the screens mapping functional related genes.

For two paralogs to have redundant or partially redundant function in the cell, they must be expressed in the same cells and at the same time. Thus, when generating such hypotheses, it can be very helpful to compare expression patterns between paralogs. To facilitate this, we integrated tissue-specific RNAseq data from publicly available resources, for example, the GTEx portal for human genes and various modENCODE RNAseq datasets for *Drosophila*. For *Drosophila*, Pearson correlation co-efficient scores for co-expression patterns are calculated and retrieved, based on modENCODE data for specific tissues, developmental stages, and cell lines. For each paralog pair, the user has the option to view the expression levels of each gene in the paralog gene pair from various datasets side-by-side as a bar graph (Figure 2).

**Figure 2.**
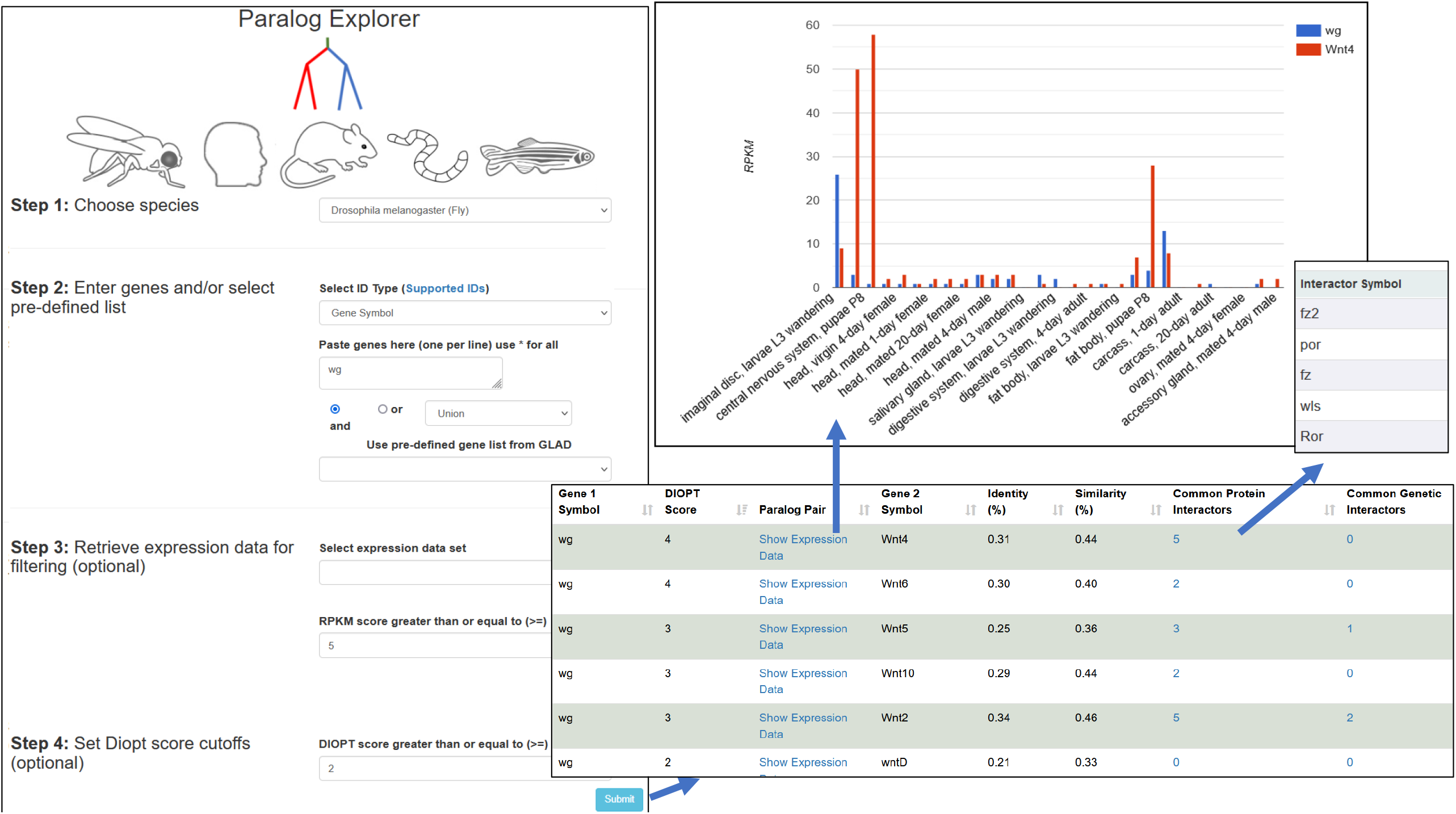
Features included in the Paralog Explore user interface. Paralog Explore is a web-based tool allowing user to select paralogs for input gene(s) along with interaction and expression data.

The user has the option to view a list of interacting partners for each gene or a list of interacting partners common to both genes (Figure 2). The choice of columns to be displayed can be customized by the user and a results table can be exported as an Excel or tab-delimited text file so that user can easily filter the list by a parameter of choice.

### Application

Paralogs exist across a very broad range of evolutionary scenarios. In the conceptually simplest cases, a gene may have a single, evolutionarily recent paralog that is highly conserved at the sequence level, and perhaps located at an adjacent location in the genome. For example, the *Drosophila* zinc finger transcription factors *gcm* and *gcm2* share 48% similarity at the amino acid level, are located just 26kb apart from one another on the second chromosome, and have been experimentally shown to retain partially redundant functions [28].

In many other cases, a gene duplication event or the duplication of part or all of the entire genome may have occurred deep in evolutionary history, creating complex gene families composed of related genes at various degrees of sequence and functional similarities. For example, the *Hox* genes [29] and most of the major developmental signaling pathways [30] underwent duplication and diversification events very early in animal evolution, leading to a scenario today where all metazoan genomes contain varying copy numbers of each member of these gene families.

In still other cases, gene families may have dramatically expanded in certain animal lineages creating exceptionally large gene families with dozens or even hundreds of members, such as the over 900 odorant receptors encoded in the mouse genome [31].

Thus, in order to be useful to researchers with various interests, Paralog Explorer should quickly, accurately, and comprehensively identify paralogs at many different scales of similarity and genomic organization, and allow the user to investigate and rank the resulting hits based on their specific research context.

We sought to test the usefulness of Paralog Explorer to identify and characterize paralogs in three typical contexts, representing a range of gene similarity and paralog number: (1) amongst recently-diverged, highly conserved pairs/triplets of conserved paralogs; (2) amongst modestly-sized gene families that duplicated and diverged early in animal evolution and have been conserved as such in modern genomes; and (3) in a large gene family containing many dozens of paralogs.

To test the usefulness of Paralog Explorer on relatively simple cases, we examined a recently published list of 25 paralog pairs or triplets in the *Drosophila* genome that are closely related and physically linked in the genome, and for which there is evidence of transcriptional co-regulation via shared enhancers [32]. For each gene pair or triplet investigated by Levo *et al*., we used Paralog Explorer to identify all predicted paralogs and ranked the results by DIOPT score. The results are presented in **Table 2**.

**Table 2.**
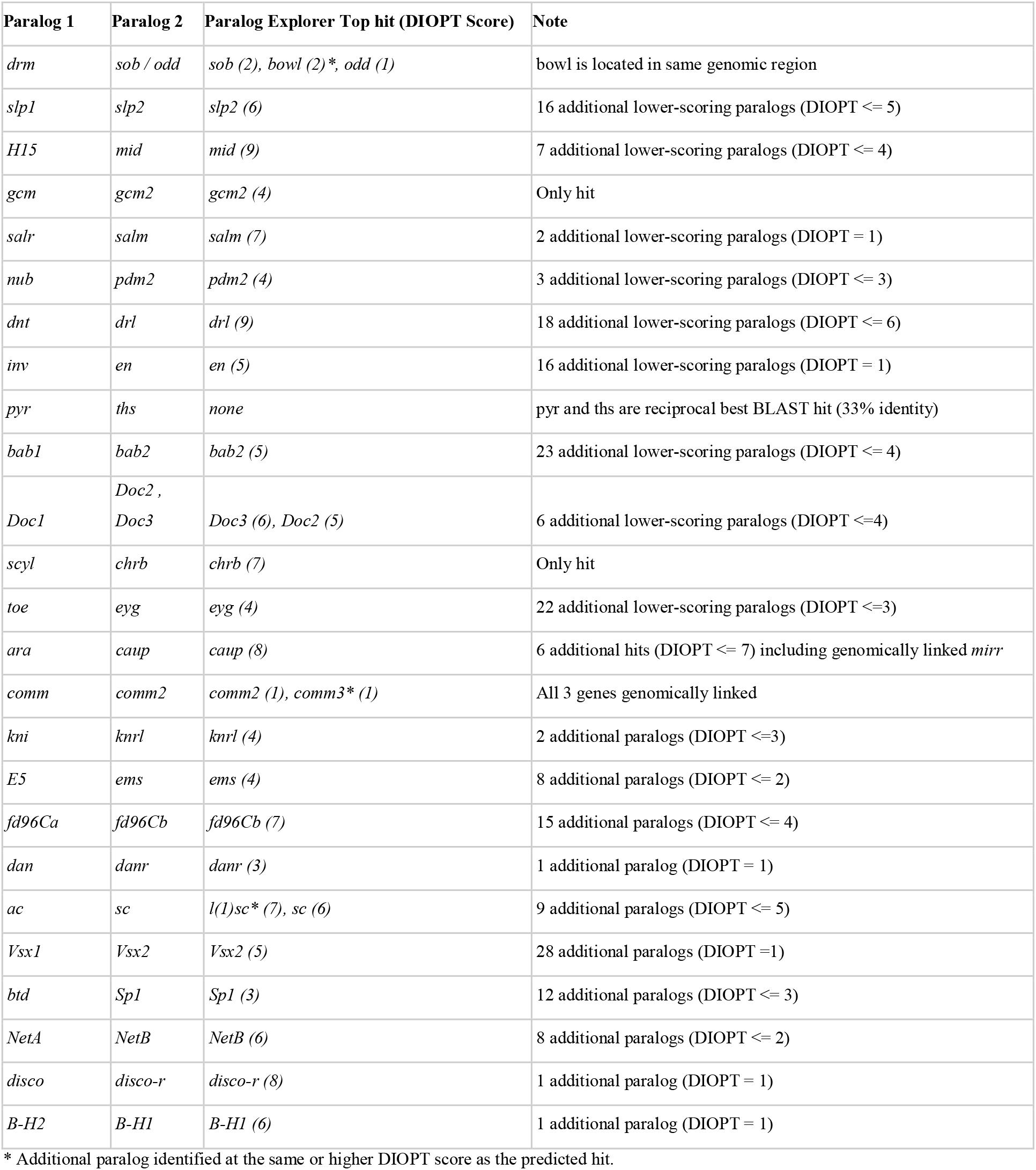
Performance of Paralog Explorer on a list of 25 curated paralog pairs/triplets from Levo *et al* (2022)

**Table 3.** Identification of multi-gene families using Paralog Explorer.

| Query Ligand | Known Paralogs | Identified via Paralog Explorer (DIOPT Score) | % of known identified |
| --- | --- | --- | --- |
| *dpp* | *gbb, scw, daw, mav, myo, Actβ* | *gbb (4), scw (4), daw (3), mav (3), myo (3), Actβ (2)* | 100 |
| *wg* | *wnt2, wnt4, wnt5, wnt6, wntD, wnt10* | *wnt2 (3), wnt4 (4), wnt5 (3), wnt6 (4), wntD (2), wnt10 (3)* | 100 |
| *dpp* | *gbb, scw, daw, mav, myo, ActB* | *gbb (4), scw (4), daw (3), mav (3), myo (3), ActB (2)* | 100 |
| *upd* | *upd2, upd3* | *upd2 (2), upd3 (1)* | 100 |
| *Pvf1* | *Pvf2, Pvf3* | *Pvf2 (1), Pvf3 (1)* | 100 |
| *spz* | *NT1 (aka spz2), spz3, spz4, spz5, spz6* | *NT1 (1), spz3 (1), spz4 (1), spz5 (2), spz6 (1)* | 100 |

In 23 of 25 cases, Paralog Explorer identified the same top-scoring paralog as was identified by manual curation [32] (Table 2), and in the remaining two instances, additional examination provided an explanation. Among the former 23 cases, Paralog Explorer returned the predicted paralog as the best-scoring DIOPT hit and allowed the viewer to quickly confirm the chromosomal location of each gene, as well as to ascertain the co-expression patterns of the gene pairs in multiple high-throughput modENCODE datasets. In several instances, Paralog Explorer identified additional high-ranking paralogs that were not listed by Levo *et al*. but which appear to be *bona fide* paralogs. For example, *bowl* is a closely related paralog of *drm, sob*, and *odd*, and is also located in the same genomic region (Table 1). Similarly, *comm3* is closely related to *comm* and *comm2*, and is located in the same genomic region (Table 2). Importantly, the existence of these additional paralogs may or may not reflect a functional conversation, but it allows researchers to systematically identify such genes for further study.

In addition to identifying the correct paralog as the top-scoring hit, Paralog Explorer also provides additional information that may be of interest. For nearly every gene query, Paralog Explorer identified a number of additional paralogs at varying degrees of similarity (Table 2). These results can be ranked by DIOPT score and/or by amino acid similarity, measures that are highly correlated with one another and serve as loose proxies for evolutionary conservation. Moreover, a user can also quickly determine whether such paralogs are physically linked in the genome and quickly access high-throughput co-expression datasets via hyperlinks.

Regarding the two cases for which Paralog Explorer did not return the same top hit as was identified via hand curation: in one case, *ac* and *sc*, Paralog Explorer identified an additional paralog, *l(1)sc*, as the top hit for *ac*, and *sc* as the second-highest hit. Thus, in this case, Paralog Explorer revealed biologically-relevant information. In the other case, *pyr* and *ths*, Paralog Explorer failed to return this pair because the current algorithms integrated by DIOPT database do not identify this pair as paralogs due to the low homology of FGF ligands [33], despite the fact that they are reciprocal best BLAST hits with one another in the *Drosophila* genome (E-value e-08, 33% amino acid identity). Because Paralog Explorer is based on the DIOPT database, this error was propagated.

Many genes belong to a complex family of multiple paralogs that duplicated and diverged at varying points during evolution, rather than as a simple pair or triplet of recently duplicated, highly-similar paralogs. For example, the TGF-β genes are a family of secreted signaling ligands that arose and diversified very early in animal evolution, and today are present in varying numbers of paralogous genes in metazoan genomes; in *Drosophila*, there are seven TGF-β genes. Phylogenetically, the seven *Drosophila* ligands fall into three sub-families: the BMP-family ligands *dpp, gbb*, and *scw*, the Activin-family ligands *daw, myo, Actβ*, and the *mav* gene which does cleanly fall into either sub-family [34]. We searched Paralog Explorer using the canonical *Drosophila* ligand *dpp*, and successfully recovered all six paralogous ligands (Figure 3). Furthermore, we noted that DIOPT scores between paralogs was generally reflective of the phylogenetic structure of the gene family [35] (Figure 3). For example, *gbb* and *scw* display the highest DIOPT score (5), and both individually score next-highest to *dpp*, resembling the taxonomic structure of these three BMP-family ligands. However, we emphasize that DIOPT scores do not directly reflect phylogenetic relationships, and can depart significantly in cases where there has been significant evolutionary change along a specific branch. For example, based on DIOPT score alone, the *Actβ* gene is most closely related to *daw* (DIOPT score = 4) and equally similar to *myo* and the other four ligands (DIOPT score = 2), whereas phylogenetic analyses reveals that *Actβ* falls into a monophyletic Activin-like group with both *daw* and *myo*, and is more closely related to both of these two paralogs than it is to the remaining four [35] (Figure 3.)

**Figure 3.**
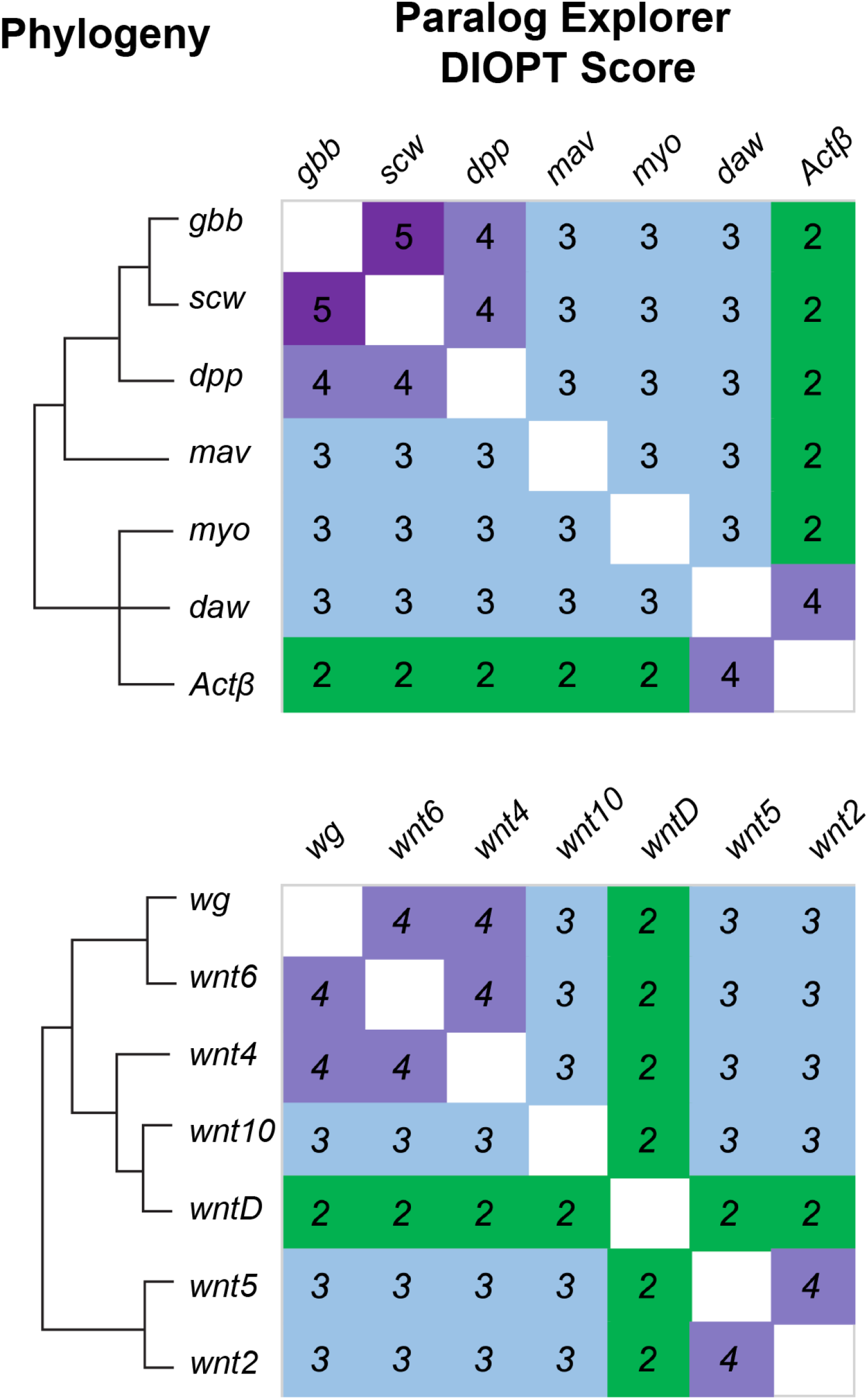
Paralog Explorer identifies known members of paralogous gene families. Using a single gene as a query, Paralog Explorer correctly identifies the complete gene family of all known TGF*β* ligands and Wnt ligands. The known gene family phylogeny is shown at left (see text for references), and a heatmap of the pairwise DIOPT scores is shown at right.

We expanded our search of gene families to include several other highly conserved signaling pathways: the seven *Drosophila* Wnt ligands, the three JAK/STAT ligands (*upd* genes), the three Pvf ligands, and the five Spatzle ligands. Each of these gene families play important roles during development, each one expanded very early in animal evolution, and each family has been expanded and/or contracted in various animal lineages. For each, we entered a single family member into Paralog Explorer, and in 100% of these examples Paralog Explorer correctly returned the entire family of related paralogs (Table 2). We note that these gene families contain a broad range of divergence amongst family members, demonstrating that Paralog Explorer is able to robustly and accurately predict the full suite of paralogs for a given gene across a wide range of evolutionary divergence and amongst complex gene families. For the Wnt family ligands, we repeated the exercise of comparing DIOPT scores to known phylogenetic relationships [36] (Figure 3). Again, dominant phylogenetic patterns of sequence conservation were reflected by DIOPT scores, while not precisely mirroring the known phylogenetic relationships. Specifically, reciprocal DIOPT scores identified *wg, wnt6*, and *wnt4* as closely related, and a close relationship between *wnt2* and *wnt5*, while the divergent *wntD* gene stood out as distinct from all other family members, all of which is reflective of known phylogenetic patterns [36]. Importantly, as with the example of TGF-β ligands shown above, results from Paralog Explorer should not be interpreted as directly reflective of phylogenetic relationships or functional conservation, but can provide potentially helpful information to generate hypothesis about genetic similarity amongst paralogs.

We then wished to know how well Paralog Explorer performed on very large gene families. We chose the odorant receptor (*Or*) gene family, of which there are 60 members in the *Drosophila* genome, as well as one pseudogene [37]. Remarkably, using *Or1a* as our query, Paralog Explorer returned exactly 59 paralogs, only failing to return the single pseudogenic member of this family noted in [37]. Thus, in even in the case of highly expanded gene families such as the *Or* genes, Paralog Explorer correctly identifies all known paralogs.

In addition to providing users with the ability to identify paralogs for individual queries or lists, Paralog Explorer also has the potential to assist in large-scale bioinformatic analyses. To demonstrate one such use case, we compared the paralog annotation with a synthetic lethality screen using CRISPR-Cas9 dual targeting [38] in human cell lines. Out of 406 heterogenous gene pairs, 21 pairs are annotated as paralogs in Paralog Explorer. Furthermore, 9 of the 21 paralog gene pairs (43%) are scored as synthetic lethality interactors with one another in at least one cell line by the criteria of FDR <0.1while only 20 out of 385 other gene pairs (5%) scored. Paralogous gene pairs are much more likely to score in functional screens than are pairs of unrelated genes, and not surprisingly, more recent studies are focused on paralog gene pairs rather than randomly selected gene pairs [6, 8, 9]. Thus, we expect that Paralog Explorer will facilitate the experimental design of high-throughput screens and mapping of functionally related genes.

There is an important caveat to Paralog Explorer, which is likely common to all paralog prediction methods. Because hypotheses of paralogy are primarily drawn from sequence conservation, gene queries which contain individual protein domains that are highly conserved may return many putative paralogs, based on the presence of shared domains across proteins that are otherwise only distantly-related. Furthermore, the sequence length of these conserved domains will impact paralog predictions, such that longer domains are more likely to score higher while relatively short domains may not reach the threshold to score via DIOPT. While the presence of shared domains across proteins may in fact reflect a true evolutionary history of gene duplication, from a practical standpoint it can lead to complex results that require sophisticated manual analysis.

As one example, we examined the *Drosophila* Hox genes, which are transcription factors that include the highly conserved homeodomain, a ∼60aa domain that is widespread across many transcription factors [29]. Inputting the anterior-most *Drosophila Hox* gene *lab* as a query, Paralog Explorer returns 23 predicted paralogs. These include the other Hox genes themselves (*pb, Dfd, Scr, Antp, Ubx, Abd-A, Abd-B*), as well as the paralogous homeodomain proteins *bcd, zen*, and *zen*-2 that are located nearby in the genome, all at a DIOPT score of 2. In addition, however, Paralog Explorer returns as its highest hit the homeobox gene *ro* (DIOPT score 3), which is not considered a Hox gene, as well as 12 additional genes, all but one of which are known homeobox-containing genes. Importantly this list is not comprehensive of all homeobox-containing genes. FlyBase identifies a total of 102 homeobox genes in the *Drosophila* genome, indicating that this hit list includes those that reach some similarity threshold based on DIOPT scores. Thus, for genes that contain specific highly conserved domains found in many genes, users should carefully analyze the results when forming hypotheses about paralogy.

## Discussion

Paralog Explorer is a tool which allows users to quickly and reliably identify paralogs of any gene(s) of interest, as well as relevant measures of their similarity, genomic location, co-expression patterns, genetic and protein interactions, and GO terms. It is important to note that identifying two or more genes as paralogs is a hypothesis about their evolutionary history – i.e. that they arose via gene duplication – rather than about molecular function and or whether they may be functionally redundant. Thus, we designed Paralog Explorer to be an inclusive and flexible search tool that will allow researchers with diverse interests to generate hypotheses about paralogous genes.

We note that Paralog Explorer is built based on predictions in DIOPT, which may miss certain paralogs that are validated by experimental data or published literature. As DIOPT is updated and improved via user-submitted data, Paralog Explorer will need to be updated accordingly.

## Acknowledgements

We would like to thank the members of the Perrimon laboratory, the FlyBase consortium, the Drosophila RNAi Screening Center (DRSC), and the Transgenic RNAi Project (TRiP) for helpful suggestions. This work was supported by NIH NIGMS P41 GM132087. N.P. is an investigator of Howard Hughes Medical Institute.

## Availability

The online resource is available without restriction at

https://www.flyrnai.org/tools/paralogs/web/.

## References

1. Ewen-Campen, B., et al., Accessing the Phenotype Gap: Enabling Systematic Investigation of Paralog Functional Complexity with CRISPR. Dev Cell, 2017. 43(1): p. 6–9.

2. Gu, Z., et al., Role of duplicate genes in genetic robustness against null mutations. Nature, 2003. 421(6918): p. 63–6.

3. De Kegel, B. and C.J. Ryan, Paralog buffering contributes to the variable essentiality of genes in cancer cell lines. PLoS Genet, 2019. 15(10): p. e1008466.

4. Ohno, S., Evolution by Gene Duplication. 1970: Springer Berlin, Heidelberg. 798812.

5. Ibn-Salem, J., E.M. Muro, and M.A. Andrade-Navarro, Co-regulation of paralog genes in the three-dimensional chromatin architecture. Nucleic Acids Res, 2017. 45(1): p. 81–91.

6. Chow, R.D., et al., In vivo profiling of metastatic double knockouts through CRISPR-Cpf1 screens. Nat Methods, 2019. 16(5): p. 405–408.

7. Han, K., et al., Synergistic drug combinations for cancer identified in a CRISPR screen for pairwise genetic interactions. Nat Biotechnol, 2017. 35(5): p. 463–474.

8. Ito, T., et al., Paralog knockout profiling identifies DUSP4 and DUSP6 as a digenic dependence in MAPK pathway-driven cancers. Nat Genet, 2021. 53(12): p. 1664–1672.

9. Thompson, N.A., et al., Combinatorial CRISPR screen identifies fitness effects of gene paralogues. Nat Commun, 2021. 12(1): p. 1302.

10. Ewen-Campen, B., et al., No Evidence that Wnt Ligands Are Required for Planar Cell Polarity in Drosophila. Cell Rep, 2020. 32(10): p. 108121.

11. Guo, L.Y., et al., Multiplexed genome regulation in vivo with hyper-efficient Cas12a. Nat Cell Biol, 2022. 24(4): p. 590–600.

12. Parvez, S., et al., MIC-Drop: A platform for large-scale in vivo CRISPR screens. Science, 2021. 373(6559): p. 1146–1151.

13. Port, F. and S.L. Bullock, Augmenting CRISPR applications in Drosophila with tRNA-flanked sgRNAs. Nat Methods, 2016. 13(10): p. 852–4.

14. Port, F., M. Starostecka, and M. Boutros, Multiplexed conditional genome editing with Cas12a in Drosophila. Proc Natl Acad Sci U S A, 2020. 117(37): p. 22890–22899.

15. Hu, Y., et al., An integrative approach to ortholog prediction for disease-focused and other functional studies. BMC Bioinformatics, 2011. 12: p. 357.

16. Hu, Y., et al., Molecular Interaction Search Tool (MIST): an integrated resource for mining gene and protein interaction data. Nucleic Acids Res, 2018. 46(D1): p. D567–D574.

17. Boley, N., et al., Navigating and mining modENCODE data. Methods, 2014. 68(1): p. 38–47.

18. Brown, J.B., et al., Diversity and dynamics of the Drosophila transcriptome. Nature, 2014. 512(7515): p. 393–9.

19. Graveley, B.R., et al., The developmental transcriptome of Drosophila melanogaster. Nature, 2011. 471(7339): p. 473–9.

20. mod, E.C., et al., Identification of functional elements and regulatory circuits by Drosophila modENCODE. Science, 2010. 330(6012): p. 1787–97.

21. Consortium, G.T., The Genotype-Tissue Expression (GTEx) project. Nat Genet, 2013. 45(6): p. 580–5.

22. Hu, Y., et al., GLAD: an Online Database of Gene List Annotation for Drosophila. J Genomics, 2015. 3: p. 75–81.

23. Ashburner, M., et al., Gene ontology: tool for the unification of biology. The Gene Ontology Consortium. Nat Genet, 2000. 25(1): p. 25–9.

24. Gene Ontology, C., The Gene Ontology resource: enriching a GOld mine. Nucleic Acids Res, 2021. 49(D1): p. D325–D334.

25. Gramates, L.S., et al., FlyBase: a guided tour of highlighted features. Genetics, 2022. 220(4).

26. Koonin, E.V., Orthologs, tparalogs, and evolutionary genomics. Annu Rev Genet, 2005. 39: p. 309–38.

27. Pereira-Leal, J.B., et al., Evolution of protein complexes by duplication of homomeric interactions. Genome Biol, 2007. 8(4): p. R51.

28. Alfonso, T.B. and B.W. Jones, gcm2 promotes glial cell differentiation and is required with glial cells missing for macrophage development in Drosophila. Dev Biol, 2002. 248(2): p. 369–83.

29. Singh, N.P. and R. Krumlauf, Diversification and Functional Evolution of HOX Proteins. Front Cell Dev Biol, 2022. 10: p. 798812.

30. Pires-daSilva, A. and R.J. Sommer, The evolution of signalling pathways in animal development. Nat Rev Genet, 2003. 4(1): p. 39–49.

31. Godfrey, P.A., B. Malnic, and L.B. Buck, The mouse olfactory receptor gene family. Proc Natl Acad Sci U S A, 2004. 101(7): p. 2156–61.

32. Levo, M., et al., Transcriptional coupling of distant regulatory genes in living embryos. Nature, 2022. 605(7911): p. 754–760.

33. Stathopoulos, A., et al., pyramus and thisbe: FGF genes that pattern the mesoderm of Drosophila embryos. Genes Dev, 2004. 18(6): p. 687–99.

34. Upadhyay, A., et al., TGF-beta Family Signaling in Drosophila. Cold Spring Harb Perspect Biol, 2017. 9(9).

35. Van der Zee, M., R.N. da Fonseca, and S. Roth, TGFbeta signaling in Tribolium: vertebrate-like components in a beetle. Dev Genes Evol, 2008. 218(3-4): p. 203–13.

36. Janssen, R., et al., Conservation, loss, and redeployment of Wnt ligands in protostomes: implications for understanding the evolution of segment formation. BMC Evol Biol, 2010. 10: p. 374.

37. Guo, S. and J. Kim, Molecular evolution of Drosophila odorant receptor genes. Mol Biol Evol, 2007. 24(5): p. 1198–207.

38. Najm, F.J., et al., Orthologous CRISPR-Cas9 enzymes for combinatorial genetic screens. Nat Biotechnol, 2018. 36(2): p. 179–189.

